# Sensory Detection of Aversive and Appetitive Stimuli by CRF neurons

**DOI:** 10.1101/209148

**Authors:** Jineun Kim, Yi-Ya Fang, Shreelatha Bhat, Anna Shin, Koichi Hashikawa, Daesoo Kim, Jong-woo Sohn, Dayu Lin, Greg S. B. Suh

**Affiliations:** Department of Biological Sciences, Korea Advanced Institute of Science and Technology, Daejeon 34141, Republic of Korea; Skirball Institute of Biomolecular Medicine, New York University School of Medicine, New York, NY 10016, USA; Neuroscience Institute, New York University School of Medicine, New York, NY 10016, USA; Department of Cell Biology, New York University School of Medicine, New York, NY 10016 USA; Department of Psychiatry, New York University School of Medicine, New York, NY 10016, USA

## Abstract

Corticotropin-releasing factor (CRF), which is released from the paraventricular nucleus (PVN) of the hypothalamus, is essential for mediating stress response by activating the hypothalamic-pituitary-adrenal (HPA) axis. CRF-releasing PVN neurons receive inputs from multiple brain regions that convey stressful events, but their rapid neuronal dynamics on the timescale of behavior remain unknown. Here, our optical recordings of PVN CRF neuronal activity in freely behaving mice revealed that PVN CRF neurons are activated immediately by a range of stress-inducing aversive stimuli, including predator odor. By contrast, CRF neuronal activity starts to drop within a second of exposure to appetitive stimuli, such as food. These findings suggest that PVN CRF neurons receive real-time information about aversive and appetitive stimuli. Their role in mediating the sensory detection and regulating the neuroendocrine axis resembles those of AgRP/NPY neurons and SFO neurons that regulate neurophysiology responses and mediate sensory detection of food and water.

---

A population of neuroendocrine neurons in the paraventricular nucleus (PVN) of the hypothalamus secretes corticotropin-releasing factor (CRF) into the circulation during exposure to stressor^1, 2^. CRF, in turn, triggers the release of adrenocorticotropic hormone (ACTH) from the anterior pituitary gland to induce the secretion of glucocorticoids (GCs) from the adrenal cortex, which comprises the final effector along the hypothalamic-pituitary-adrenal (HPA) axis^1, 2^. This gradual two-step hormonal mechanism of HPA activation adjusts the neuroendocrine responses to stress. It does not, however, explain how PVN CRF neurons are influenced by rapid changes in the regulatory activities responsible for the behavioral changes that occur in response to acute stress. As such, the rapid response to display defensive behavior would require a new pathway.

To understand the response of PVN CRF neurons to stressful stimuli in awake, freely behaving mice, we used fiber photometry, *in vivo* calcium imaging technique that measures the total fluorescence of a calcium reporter expressed in a population of neurons. We virally expressed a Cre-dependent GCaMP6s calcium indicator in CRF-ires-Cre mice, and recorded changes in GCaMP6s fluorescence signals through a 400µm optic fiber placed above the PVN (**Fig. 1a**). The resulting trace represents the integrated activity of PVN CRF neurons. Histological analysis revealed that approximately 95% of GCaMP6-expressing cells overlap with CRF-positive cells in the PVN (**Fig. 1a**).

We first investigated changes in GCaMP fluorescence intensity in PVN-CRF^GCaMP6^ mice that were subjected to various stress-inducing conditions. During a forced swimming test, which increased *c-fos* expression in PVN CRF neurons significantly^3^, *in vivo* recordings revealed a substantial increase in GCaMP intensity when the mice were dropped into the water, which rose continuously until the mice were removed from the water (**Fig. 1b, c**). The GCaMP intensity remained high when the mice were returned to their home cage but returned to pre-test baseline levels over 20 minutes later (**Fig. 1c**). During tail-restraint test, we also observed a rapid increase in GCaMP intensity in PVN CRF neurons when the mice were lifted by hand (**Fig. 1d, e and Supplementary Fig. 1a, c**). The GCaMP signals declined sharply, however, when the mice were released. After several rounds of tail-restraint tests, the baseline GCaMP fluorescence gradually increased (**Fig. 1d**).

PVN CRF neuronal activity also increased in mice exposed to other exteroceptive stressors, such as a looming stimulus or predator odor. The looming stimulus consisted of a pigeon-sized object moving above the mice to mimic a flying predator of these mice (**Fig. 1f, g and Supplementary Fig. 1b, d**). These signals did not increase significantly, however, when the object was ‘flying’ lateral to the mice (**Supplementary Fig. 2a**). To investigate this effect further, we compared the response of PVN CRF neurons when the mice were exposed to an illuminating looming disk above, along the side or below a transparent cage in which the mice were placed. The disk looming above the cage, which triggered robust defensive behavior in mice^4^, resulted in the strongest activation of PVN CRF neurons (**Supplementary Fig. 2b**).

Aversive olfactory stimuli also stimulated the activity of PVN CRF neurons. Exposure to a predator odor, 2,3,5-trimethyl-3-thiazoline (TMT), triggered bursts of flight and freezing responses in mice^5^. We found a significant increase in GCaMP signals in these neurons when the mice displayed the defensive responses to TMT (**Fig. 1h, i**). These stress-inducing aversive stimuli of different sensory modalities rapidly stimulated the activity of PVN CRF neurons (**Fig. 1j**).

We next investigated the response of PVN CRF neurons when the mice were subjected to an interoceptive stressor, food deprivation, followed by food consumption. Following 22 hours of food deprivation, GCaMP signals were increased in the PVN CRF neurons of these mice (**Fig. 2** and Data not shown). This is consistent with the increase in *c-fos* expression detected in PVN CRF neurons during prolonged periods of food deprivation (**Supplementary Fig. 3**). Control mice with PVN CRF neurons expressing GFP showed no change in fluorescent intensity during these stress-inducing conditions, suggesting that the observed changes in ∆F/F was not due to a movement artifact (**Supplementary Fig. 4**).

While it is unexpected that aversive stimuli rapidly stimulated the activity of PVN CRF neurons, their role in activating the HPA axis has been well established. It is completely unknown, however, whether PVN CRF neurons respond to attractive stimuli. To investigate the response of these neurons to such stimuli, we measured changes in GCaMP intensity in PVN CRF neurons in response to food. We first introduced a control pellet-sized neutral object into their home cage, followed by a normal food chow pellet. (**Fig. 2a**). The intensity of the GCaMP signal in these neurons was initially dropped slightly when some mice inspected the control object but quickly returned to its previous level (**Fig. 2a** and see arrow). By contrast, PVN CRF neuronal activity dropped precipitously when the mice consumed chow pellet (**Fig. 2a-c**). Moreover, the fluctuation in the GCaMP signals was substantially suppressed (**Fig. 2a, c**). Intriguingly, GCaMP signals started to drop even before the mice took their first bite of food (**Fig. 2c-inset**). To determine whether the sensory cue of food is sufficient to reduce the activity of PVN CRF neurons, we provided chow pellet in a mesh-covered cup to prevent physical contact with it while allowing the mice to see it and smell it (**Fig. 2d**). In these mice, GCaMP signals were significantly suppressed initially, but were soon increased to higher levels (**Fig. 2d-f** and see arrowheads). The kinetics of the GCaMP suppression in these mice was initially indistinguishable from those in mice freely consuming chow pellet (**Fig. 2g**; τ=35.8±6.3s for chow/cup and τ=32.1±12s for chow in fasted state; τ=664.1±70s for chow in fed state). In contrast to food-deprived mice, the food-induced suppression of GCaMP activity was not observed in fed mice (**Fig. 2b, e**).

We then determined the effect of attractive odorants on PVN CRF neuronal activity. Exposure to highly palatable peanut oil^6^ resulted in a small, yet significant decrease in the intensity of GCaMP signals in these neurons (**Supplementary Fig. 5**). Similarly, when the mice were exposed to 2-phenylethanol (2-PE)^6^, which elicits attraction behavior in mice, the activity of PVN CRF neurons also decreased, but the effect was not significant.

We next sought to measure the activity of PVN CRF neurons during social interactions with intruders. We introduced a pup or a male aggressor into the home cage of a mouse in which GCaMP activity in PVN CRF neurons was continuously recorded. PVN CRF neuronal activity in a female mouse regardless of whether she is lactating or naïve was robustly suppressed when a pup was introduced to her cage and was continuously suppressed during the period when the female interacts with the pup (**Fig. 3a-c**). By contrast, when a recorded male was attacked by an aggressive male intruder, we observed rapid, dramatic increases in GCaMP signals in PVN CRF neurons during attack (**Fig. 3d-f**). Even after the end of the episode, the GCaMP activity did not return to the original baseline (**Fig. 3d**). When male-male interactions did not involve aggression, we did not observe a noticeable change in PVN CRF neuronal activity (**Supplementary Fig. 6**). When the recorded mouse was presented with a fake animal, the activity of PVN CRF neurons did not change (**Fig 3g, h**).

Our fiber photometry recordings of PVN CRF neurons in freely behaving mice suggest that these neurons mediate the sensory detection of a wide range of stressors of various modalities. The speed of their response suggests that it is probably mediated by neuronal rather than hormonal input. One function of this neuronal input would be to inform PVN CRF neurons of the presence of a stressor. To do so, it would have to do more than merely stimulate a neuroendocrine response through a hormonal cascade; this could take tens of minutes to occur. This dual function closely resembles that of AgRP hypothalamic neurons that respond to energy homeostasis but also mediates the sensory detection of food^7^, and SFO neurons that respond to fluid homeostasis, but also mediate the sensory detection of water^8^. Each of these may represent a straightforward means of quickly pairing the physiological needs of the body with the anticipation of finding food or water in the environment. The speed with which both aversive and attractive stimuli are detected by PVN CRF neurons may serve as an effective way to reset their ongoing regulation of the neuroendocrine HPA axis.

Furthermore, the speed with which mice display defensive behaviors following stress are at odds with the slowness of the neuroendocrine response to stress. Thus, it is likely that the immediate activation of PVN CRF neurons is necessary to trigger rapid defensive behavior. Indeed, acute photostimulation of PVN CRF neurons in mice was followed by behavioral responses similar to those that are typically observed at the onset of acute stress, such as grooming^9, 10^. The observation that CRF knockout mice exhibited normal grooming behavior following stress, however, suggests that this stress-induced behavior is mediated by a pathway independent from the HPA axis^11^. Further study may allow us to identify a distinct pathway through which PVN CRF neurons mediate rapid defensive behaviors in response to stress and, possibly, facilitate our discovery of a key mechanism underlying the response to acute stress.

Our *in vivo* recordings revealed unexpectedly that the activity of PVN CRF neurons is rapidly and potently suppressed by appetitive stimuli. Previously, quantitative immune-electron microscopy studies illustrated that approximately half of all synapses onto PVN CRF neurons are GABAergic and the other half are glutamatergic^12^. While the glutamatergic input potentiates PVN CRF neuronal activity, the GABAergic synaptic transmission onto PVN CRF neurons was proposed to modulate the glutamate-induced activation of HPA axis^13, 14^. The substantive GABAergic input, however, may allow PVN CRF neurons to encode both aversive and attractive signals to regulate appropriate behavioral responses.

## METHODS

### Mice

All animal experiments were performed according to protocols approved by NYU and KAIST IACUC protocols for the care and use of laboratory animals. We used CRF-ires-Cre mice (B6(Cg)-*Crh*^*tm1(cre)Zjh*^/J; Jackson Laboratory), both female and male (8-16weeks old) housed individually under a 12-hr light/dark cycle (7 am to 7 pm light), with food and water available *ad libitum* unless specified.

### Stereotaxic surgery

120nl of AAV1-CAG-Flex-GCaMP6s virus (University of Pennsylvania Vector Core) was stereotaxically injected unilaterally into the PVN (Bregma ML:0.2 mm, AP: −0.75 mm, DV: −4.7 mm from the brain skull) of CRF-ires-Cre mice using a Nanoject III (Drummond). In the same surgery, a mono fiber-optic ferrule (5 mm, 400µm CD, 0.48 NA; Doric lenses) was implanted above the PVN (-4.3 mm DV) and sealed with dental cement (C&B Metabond, S380). Animals received intraperitoneal injection of 5mg/kg ketoprofen after the surgery and were singly housed to recover for 3-4 weeks before testing.

### Fiber photometry recording

The fiber photometry setup was constructed as previously described^1,2^. Briefly, a 400 Hz sinusoidal 473 nm blue LED light (30 μW) (LED light: M470F1; LED driver: LEDD1B, Thorlabs) was band-pass filtered (passing band: 472 ± 15 nm, Semrock, FF02-472/30-25) and used to excite GCaMP6s fluorescence. A dichroic mirror was used to reflect 473 nm and pass the emission light from the activated GCaMP6s fluorescence. Bandpass filtered light (passing band: 534 ± 25 nm, Semrock, FF01-535/50) was detected by a femto-watt silicon photodetector (Newport, 2151) and recorded as digitized signal using a real-time processor (RP2.1, TDT). The real-time signal was time-locked following the same time frame from the video recording system using StreamPix such that the acquired signals and the monitored episodic behaviors of the animals can be synchronized.

The normalized GCaMP fluorescence signals (Fn) were calculated from dividing the raw GCaMP signals by the mean fluorescence signals from −100 sec to ~-5 sec before the stimulus presentation.

For the peri-event time histograms (PETHs), the onsets of each event were aligned to the time zero and the signals were normalized as (F – F_baseline_)/F_baseline_ × 100 (ΔF/F %). F_baseline_ was the mean of GCaMP signals for the window before the time zero. F is the mean of GCaMP6 signals for the window after the time zero.

For the population plots in Figure 1c, i, Figure 2 and Figure 3c, F_baseline_ was calculated from −100 sec to ~-5 sec before the stimulus introduction to exclude the increased signals due to the placement of stimuli in the box.

**Figure 1.**
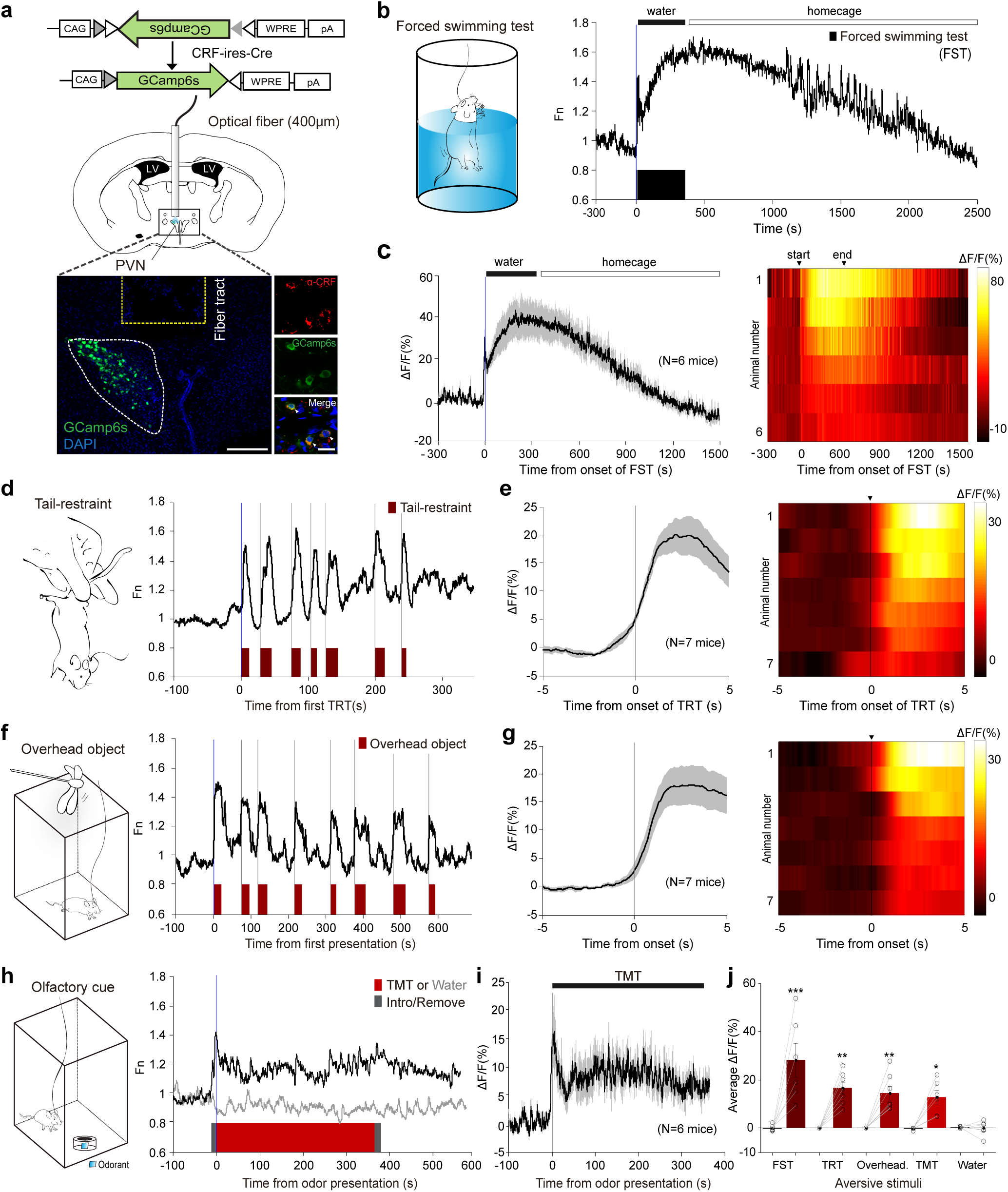
Acute increase in PVN CRF neuronal activity by naturally occurring aversive stimuli. **(a)** Viral construct and implantation scheme for fiber-photometry on PVN-CRF neurons. Bottom left – a representative image validates GCaMP6s expression in CRF neurons and optical fiber tract above the PVN. Scale bar: 200µm; Bottom right – images depict the overlap between GCaMP6s expressing cells (green) and anti-CRF positive cells (red). Scale bar: 20µm. (**b**) Schematic for forced-swimming test (FST) and a representative trace illustrating an increase of PVN-CRF^GCaMP6^ signal during FST (black bar) and sustained activity while in back to home cage (white bar). (**c**) Plot (left) and heat map (right) across animals aligned to the start and end of FST, and the following rest in home cage. Black bold line and grey shadow in this and following figures indicate mean and s.e.m., respectively. (**d**) Schematic for tail-restraint test (TRT) and a representative trace showing increases of PVN-CRF^GCaMP6^ signal during TRT (red bars). **(e)** PETH plot (left) and heat map (right) across animals aligned to the start of TRT. **(f)** Schematics for presenting an overhead object and a representative trace showing increases of PVN-CRF^GCaMP6^ signal during the presentation (red bars). **(g)** PETH plot (left) and heat map (right) across animals aligned to the start of the presentation. **(h)** Schematic of odor presentation and a representative trace showing increases of PVN-CRF^GCaMP6^ signal during TMT exposure (red bar). The periods during which mice were introduced or removed were depicted in grey bars. **(i)** Plot across animals aligned to the start and end of TMT exposure. 6 out of 9 mice that displayed a defensive response showed an increase in GCaMP signal. **(j)** Comparison of average ΔF/F of PVN-CRF^GCaMP6^ in response to different aversive stimuli. Paired two-tailed student’s t-test for raw GCaMP signals before vs. during exposure. *p<0.05, **p<0.01. Data are presented as mean ± s.e.m.

**Figure 2.**
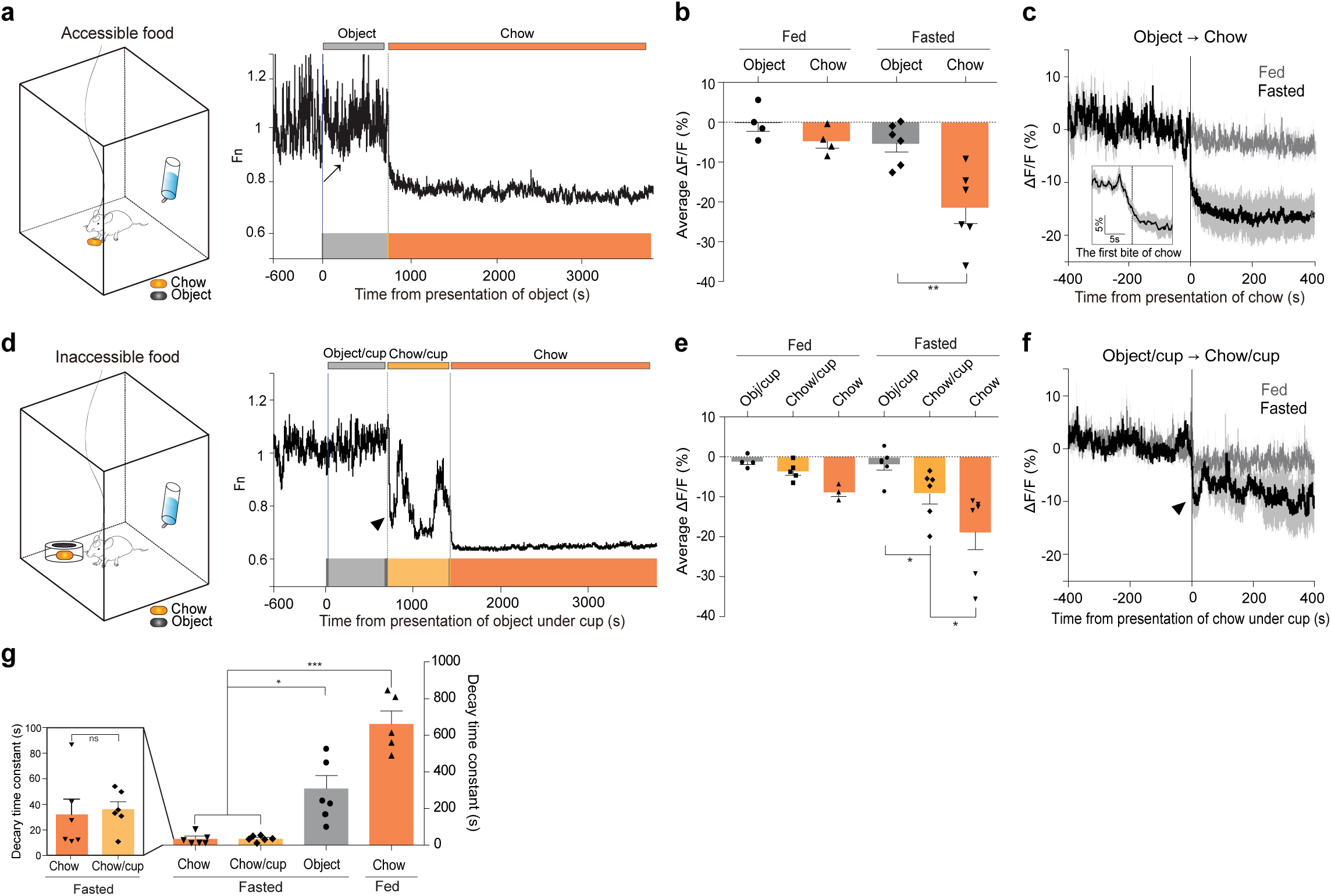
Suppression of PVN CRF neuronal activity by appetitive cues. **(a)** Schematic for freely-accessible chow presentation in a chamber and a representative trace showing PVN-CRF^GCaMP6^ signal during exposure to different cues. 22-hr fasted mice were freely exposed to non-food object (grey bar) followed by chow pellet (orang bar). Arrow depicts a drop in PVN-CRF^GCaMP6^ signal when a mouse inspects non-food object. **(b)** Bar graph summarizing average ΔF/F of PVN-CRF^GCaMP6^ when exposed to non-food object and chow pellet in fed or fasted mice. **(c)** Plot across animals aligned to the introduction of chow pellet (N=6). Inset: plot across animals aligned to the first bite of chow pellet. **(d)** Schematic for inaccessible chow presentation in a chamber and a representative trace showing PVN-CRF^GCaMP6^ signal during exposure to different cues. 22-hr fasted mice were exposed to non-food object (grey bar) and then chow pellet (yellow bar) in a mesh-covered cup followed by freely accessible chow pellet (orang bar). Arrowhead depicts a drop in PVN-CRF^GCaMP6^ signal when mice inspect chow pellet in a mesh-covered cup. **(e)** Bar graph summarizing average ΔF/F of PVN-CRF^GCaMP6^ when exposed to non-food object followed by chow pellet in a mesh-covered cup. **(f)** Plot across animals aligned to the introduction of inaccessible chow pellet in a mesh-covered cup (N=6). **(g)** Bar graph illustrating the decay time constant when exposed to accessible non-food object and chow pellet, and inaccessible chow pellet in a mesh-covered cup. Left graph magnifies the two bars representing accessible chow and inaccessible chow. Paired or unpaired student’s t-test. *p<0.05, **p<0.01. Data are presented as mean ± s.e.m.

**Figure 3.**
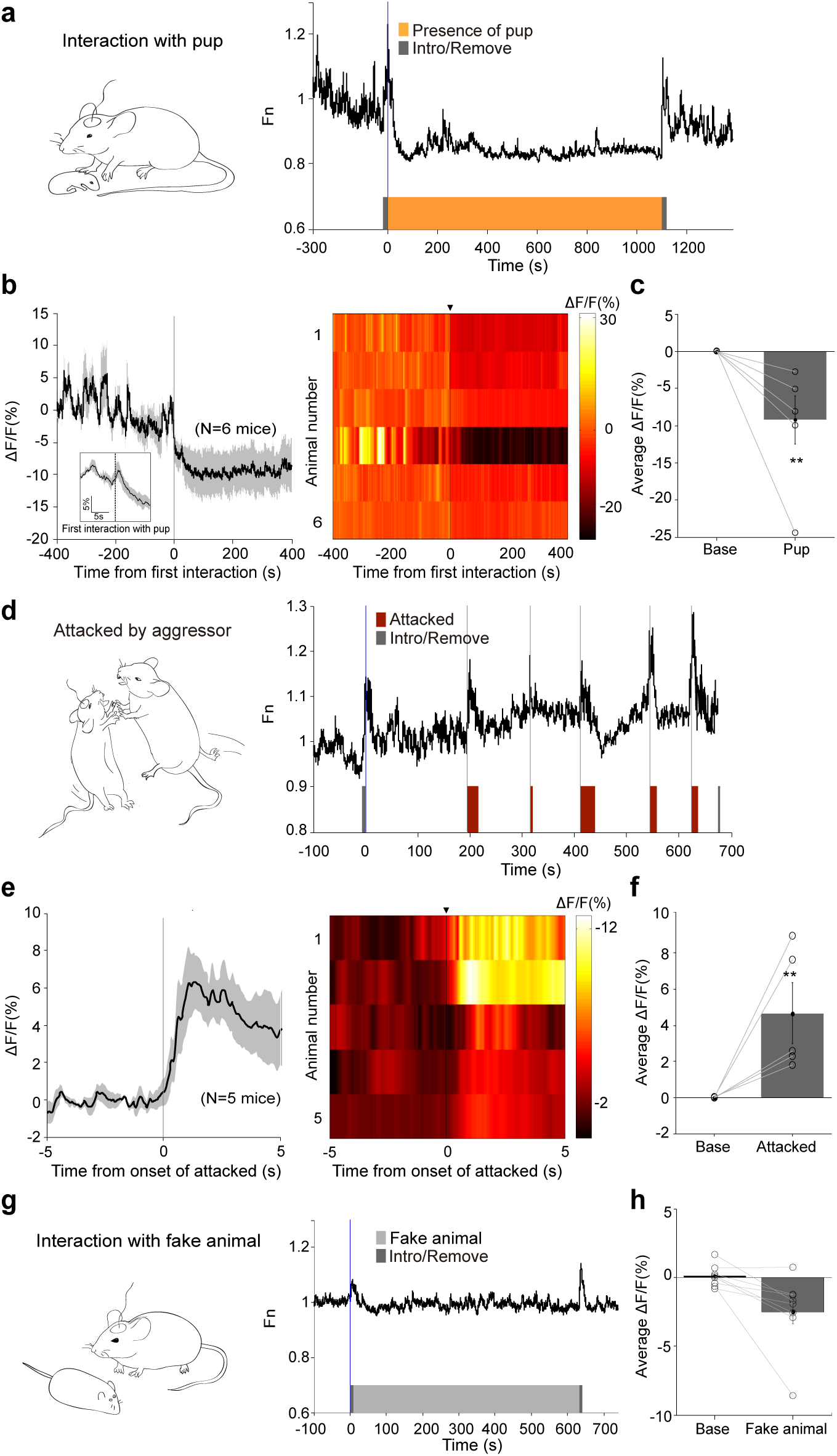
PVN CRF neuronal responses during social interaction. **(a)** Schematic for ‘interaction with pup’ and a representative trace illustrating suppression of PVN-CRF^GCaMP6^ in a female mouse during interaction with pup (orange bar). The periods during which mice were introduced or removed were depicted in grey bars. **(b)** Plot (left) and heat map (right) across animals aligned to the onset of interaction with pup (N=6 female). Inset: plot across animals aligned to the onset of interaction with pup in a 10-s time window. **(c)** Bar graph demonstrating a decrease in average ΔF/F of PVN-CRF^GCaMP6^ during interaction. **(d)** Schematic for ‘attacked by aggressor’ and a representative trace illustrating acute surge in PVN-CRF^GCaMP^ signal to the onsets of being attacked (red bars). **(e)** PETH plot (left) and heat map (right) across animals aligned to the onset of attacked (N=5 male). **(f)** Bar graph demonstrating an increase in average ΔF/F of PVN-CRF^GCaMP6^ during attacked. **(g)** Schematic for ‘interaction with fake animal’ and a representative trace during the presentation of fake animal. **(h)** Bar graph showing the average ΔF/F of PVN-CRF^GCaMP6^ during the episode Paired two-tailed student’s t-test. *p<0.05, **p<0.01. Data are presented as mean ± s.e.m.

In Figure 2g, the decay time constant during the initial phase of drop in GCaMP signal was obtained as the duration between the onset of stimulus presentation and the time at which the GCaMP6 signal reaches 63.2% of the maximal change.

### Behavior test and video recording

All experiments were performed in a custom-made behavior chamber with air-fan and under dimmed light from the start of dark cycle. Mice were habituated in the chamber for 20 mins before each experiment. Their behaviors were video-taped using an infrared camera (Basler, ace120gc) placed from the top and side of the chamber. All behaviors were further analyzed by manually annotating using custom software written in MATLAB (http://vision.ucsd.edu/~pdollar/toolbox/doc/index.html). The behavior annotation was time-locked with the recorded GCaMP6 signal and plotted as a bar plot under its trace using MATLAB.

### Forced swimming test (FST)

We used the protocol for forced swimming test that was described previously^3, 4^. Briefly, the animal was gently placed into 3 L of beaker (18 cm of height) filled with 24-25℃ of water until 12 cm. After 6 mins of forced swimming, the mice were gently dried by paper towel and placed back in their home cage. 6 mice were tested on 3 different days.

### Tail restraint test (TRT)

The tail of tested mice was chased and grabbed by hand. After grabbing, the mice were suspended in air for 6-8 sec before releasing them in their home cage. 7 mice were tested on 5 different days; each mouse was tested at least 5 times.

### Presentation of a looming cue

A large bird-like object was custom made and presented as a cue that moves above or lateral to the cage. For more systematic presentations of a visual cue, we engineered a looming shadow disk as previously reported^5^. The looming shadow disk panel was placed at the bottom, side or top of a transparent cage (25 × 25 × 30 cm) sequentially with 10 mins of interval after 15 mins of habituation. The ‘Looming disk’ was programmed as a black circle in grey background, increasing its size from 2 degree of visual angle expanding to 20 degree in 250 ms, maintained for 250 ms and presented repeatedly for 15 times with 500 ms of interval for one trial. 7 mice were tested on 5 different days; each mouse was tested at least 3 times per condition.

### Presentation of odorants

Mice were habituated for 10 mins in a chamber box (20 cm x 20 cm x 20 cm) equipped with an air-fan. A piece of 2 cm^2^ filter paper soaked with 40 μl of either water, 80 mM of peanut oil, 2-phenylethanol (2-PE), 70% ethanol, or 1:10 diluted 2,5-dihydro-2,4,5-trimethylthiazoline (TMT, Sigma Aldrich) was presented gently at the center of the chamber as described^6^. The odorants were presented for 5 mins with 15 mins of interval. 6 mice were tested on 6 different days.

### Presentation of food or food-cues

On day 1, baseline activity in *ad libitum* fed state was measured for 1 hr in a chamber (20 × 20 × 20 cm, equipped with water sipper). Following 1 hr of recording in the chamber, the tested animal was deprived of food with ad-lib water for 22 hrs in their home-cage. On day 2, the fasted animal was replaced into the behavior chamber and habituated for 15 mins. After the habituation, a small object (~1 cm^3^) was presented for 10 mins as a control and then a small pellet of food (4 g/pellet) was placed. Mice were observed for at least 45 mins before the start of food pellet consumption.

For the inaccessible food condition, mice were habituated in the chamber equipped with a meshed cup (9 cm of diameter, 3 cm of height). The object or chow pellet was placed under the meshed cup for 10 mins, followed by free access to the chow pellet. 6 mice were tested on 3 different days

### Resident-intruder social interaction

To observe social interactions and PVN CRH neuronal activity *in vivo*, different intruders were brought into the home cage of recorded mice. A pup (<2 weeks) was carefully presented into a female mouse to observe the pup-female interaction. Aggressive CD1 male intruder mice were selected as previously reported^7^. The same CD1 aggressor was introduced into the home cage of the recorded males on different days. Post-recording video annotation identified the epochs of ‘being attacked’. As a control, ‘fake animal’ – a mouse-like object that is similar size as adult B6 mice was used.

### Immunohistochemistry

Animals were anesthetized with isoflurane and cordially perfused with 30 ml of 4% PFA following 30 ml of 0.9% saline. Brains were post-fixated for 4 hrs in 4% PFA at 4℃ and then transferred to 20% and 30% sucrose in PBS serially. Brains were coronally sectioned at 50 μm using a Leica cryostat. The sectioned brains were washed with PBS for 10mins and blocked with 2% normal donkey serum in 0.2% PBST for 1 hr at RT. Primary antibodies; rabbit anti-CRF (1:250, late Wylie *Vale lab,* PBL rC68) or goat anti-cFos (1:200, Santa Cruz, sc 52-g) in the blocking solution (2% NDS, 0.2% PBST) were incubated at 4℃ for 72 hrs or 24 hrs, respectively. The sections were washed with PBS (3 × 15 mins), followed by incubation of second antibodies – donkey anti-rabbit Alexa 647 (1:500, Life Technologies, A21207) or donkey anti-goat Alexa 488 (1:500, Life Technologies, A11055) for 1 hr at RT. Sections were washed with PBS (2 × 15 mins), stained with DAPI (1:10000) for 6 mins, mounted on slides and coverslipped with DAKO mounting medium. Confocal images were captured with Zeiss LSM 780 microscope and analyzed using Image J.

### Statistical analysis

All statistical analysis done in MATLAB or Prism software. Unpaired or pair-wised comparison was done using two-tailed student’s t-test. Data are presented as mean ± s.e.m. Details of statistics done written in related figure legends.

## ACKNOWLEDGMENTS

We are grateful to Kalyani Narasimhan, Leandro Vendruscolo, and members of the Suh laboratory for critical comments on the manuscript. We thank the laboratory of late Wylie Vale for an aliquot of anti-CRF antibody. This work is supported by TJ Park Science Fellowship and KAIST Innovative Doctoral Research Fellowship to J. K., NIH grants (RO1MH101377 to D.L. and RO1DK106636 to G.S.B.S.), and KAIST Chancellor’s fund to G.S.B.S. and J.W.S.

## AUTHOR CONTRIBUTIONS

G.S.B.S., D.L., and J.K. conceived the project, designed the experiments, and interpreted the results. G.S.B.S. wrote the manuscript with D.L. and J.K. J.K. performed all the experiments with assistance from Y.F and S.B. J.K. and D.L. analyzed calcium imaging data. Y.F., K.H., and D.L. made possible for J.K. to carry out Fiber Photometry imaging experiments.

---

**Supplementary Figure 1.**
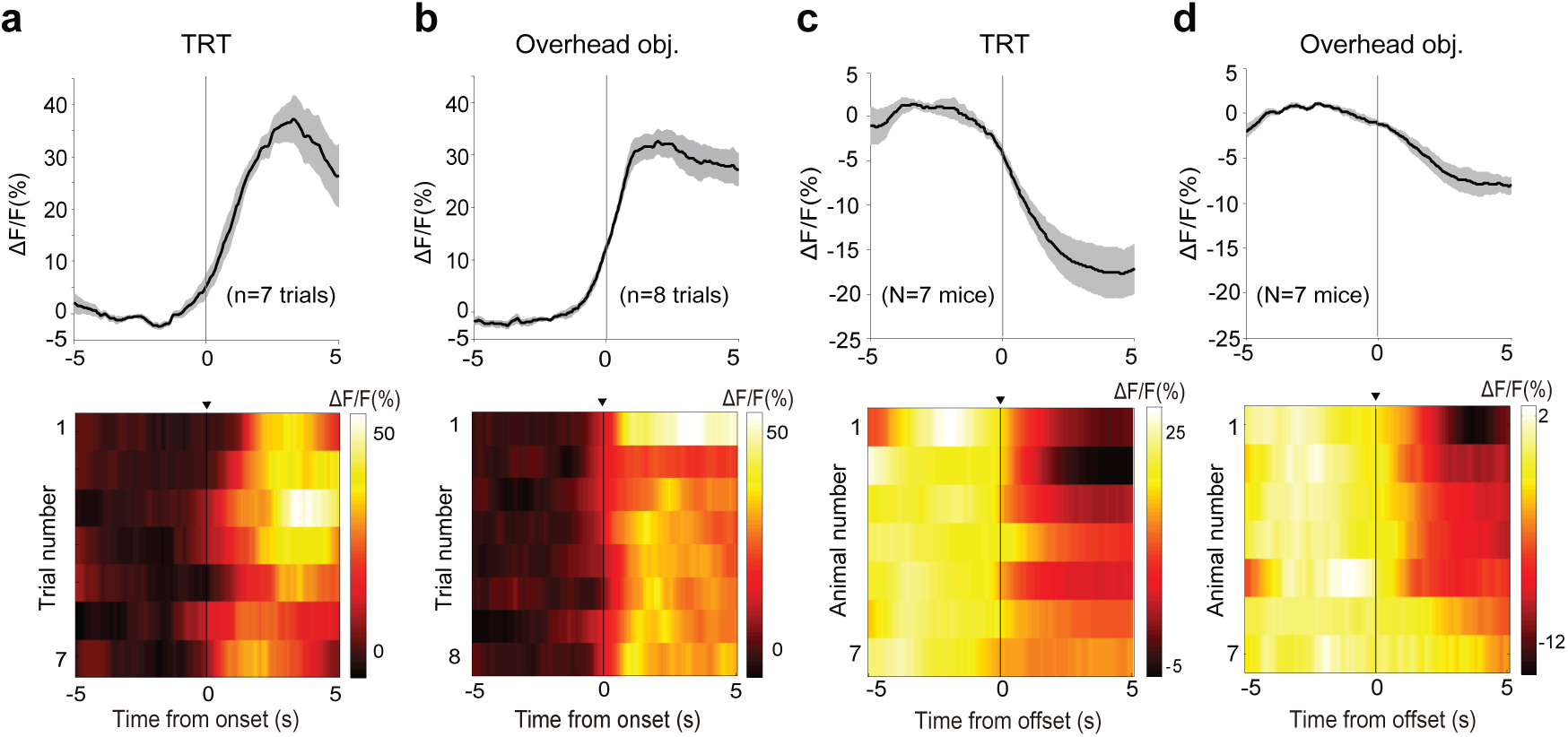
PVN-CRF^GCaMP6^ signal is increased by the onset and is decreased by the offset of exposure to aversive stimuli. **(a-b)** PETH plot (upper) and heat map (bottom) from a representative animal shown in Figure 1d (TRT) **(a)** and in Figure 1f (overhead object) **(b)**. **(c-d)** PETH plot (upper) and heat map (bottom) for population activity across animals to the offset of TRT **(c)** and the offset of overhead object **(d)**. Related to Figure 1.

**Supplementary Figure 2.**
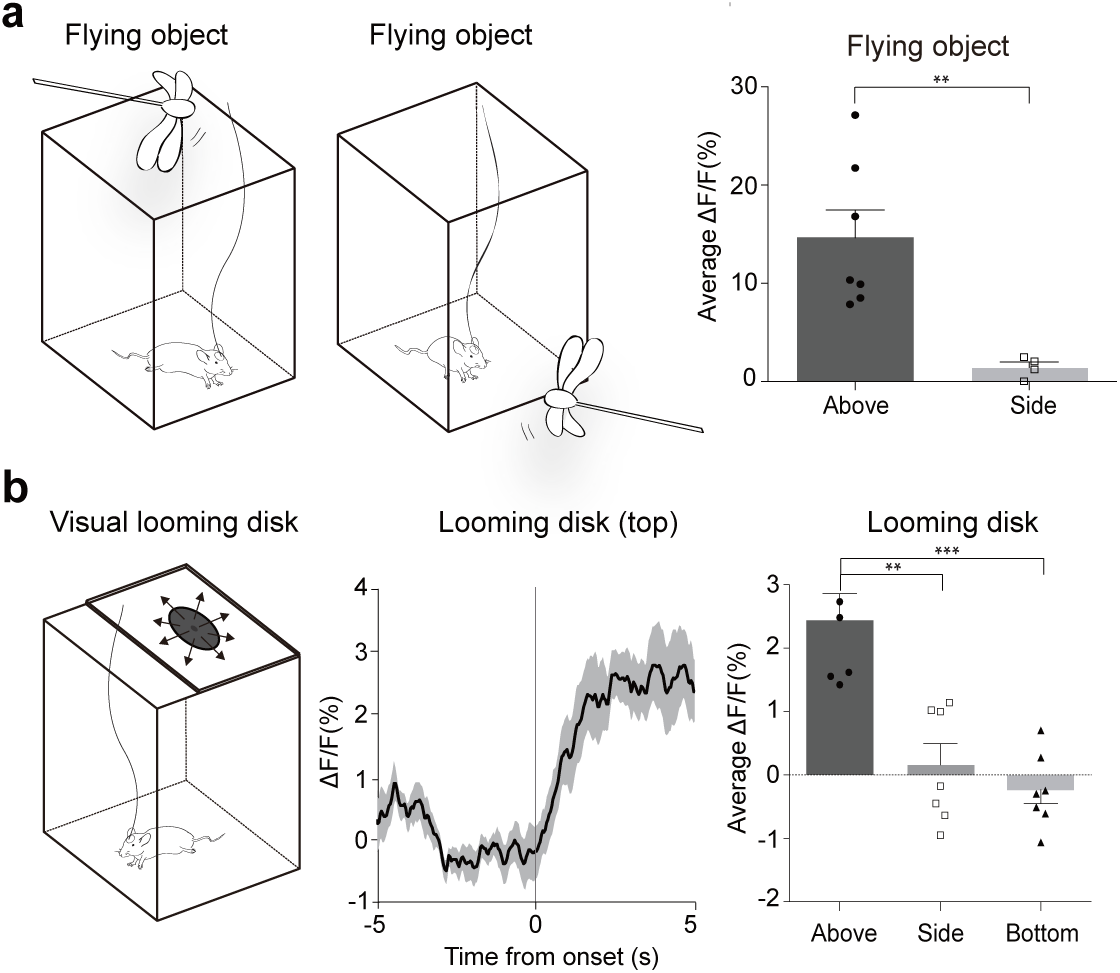
PVN-CRF^GCaMP6^ signal is increased by a ‘flying’ object presented from above, but not from side or bottom. **(a)** Schematic for a ‘flying’ object from above and side (left). Bar graph showing average ΔF/F of PVN-CRF^GCaMP6^ in mice that were exposed to the flying object from above (N=7) and side (N=4) (right). **(b)** Schematic for a looming shadow disk (left) and PETH plot for population activity across animals to the onset of the looming disk (middle). Bar graph showing average ΔF/F of PVN-CRF^GCaMP6^ (a 5-s window after time zero) in mice that were exposed to the disk from above, side or bottom (right) (N=7). **p<0.01 ***p<0.001, Data are presented as mean ± s.e.m. Related to Figure 1.

**Supplementary Figure 3.**
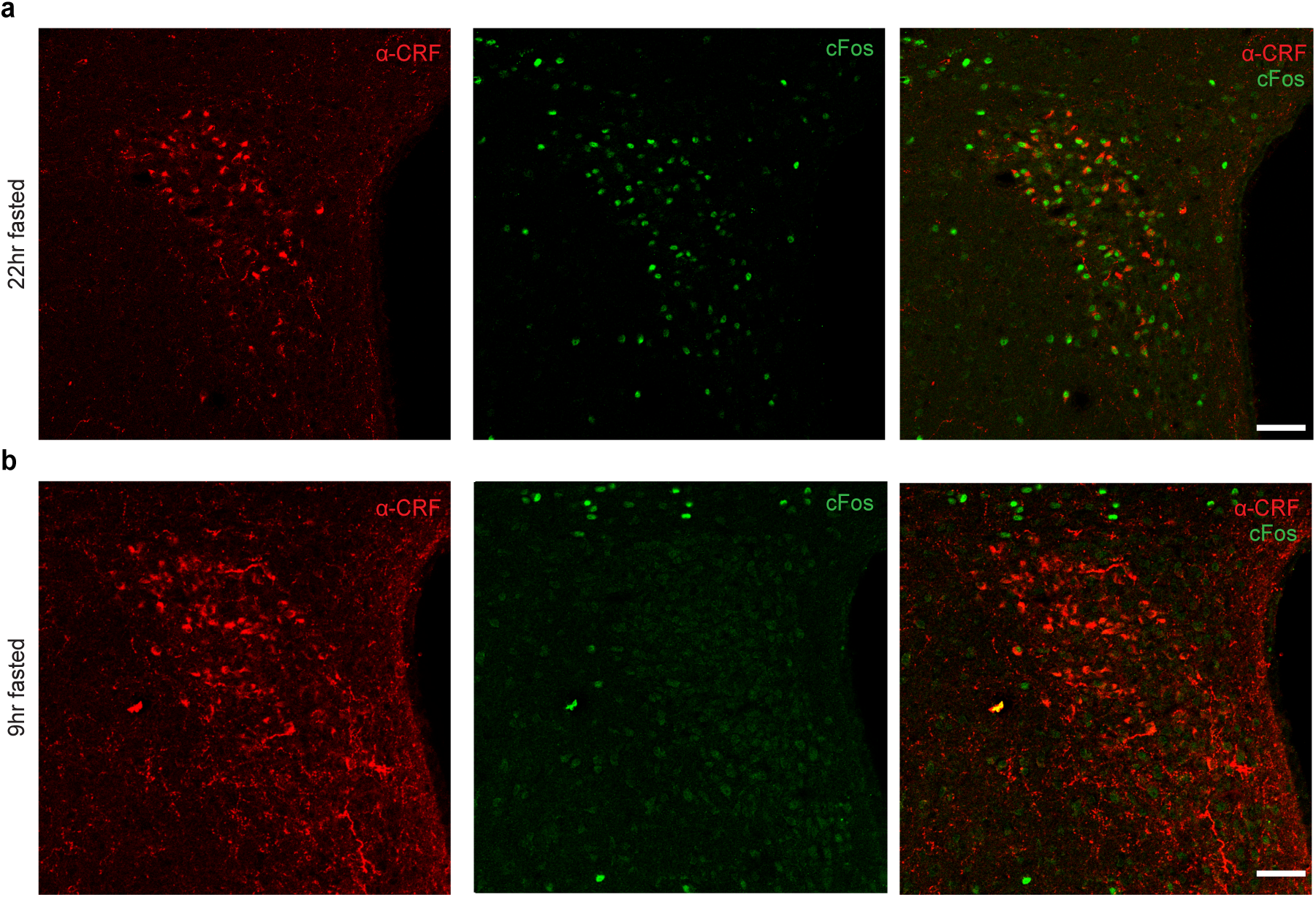
cFos induction in PVN CRF neurons after periods of starvation. **(a-b)** A representative confocal image of PVN in a mouse fasted for 22 hours **(a)** or fasted for 9 hours **(b)** immunostained with anti-CRF (red) and anti-cFos (green) antibodies. Scale bar: 50μm. Related to Figure 1.

**Supplementary Figure 4.**
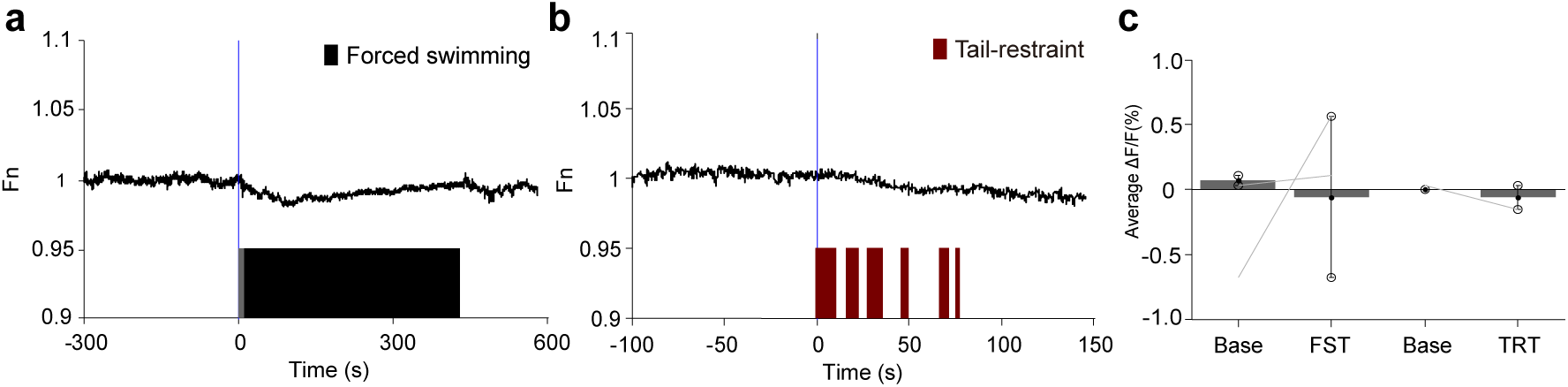
The changes in ∆F/F of PVN-CRF^GFP^ signal in response to environmental stimuli was not due to a movement artifact. **(a)** A representative trace illustrating no change in the normalized PVN-CRF^GFP^ signal during FST (N=2). **(b)** A representative trace illustrating no change in the average PVN-CRF^GFP^ signal during TRT (N=2). **(c)** Bar graph illustrating average ΔF/F of PVN-CRF^GFP^ signals during FST and TRT. Related to Figure 1.

**Supplementary Figure 5.**
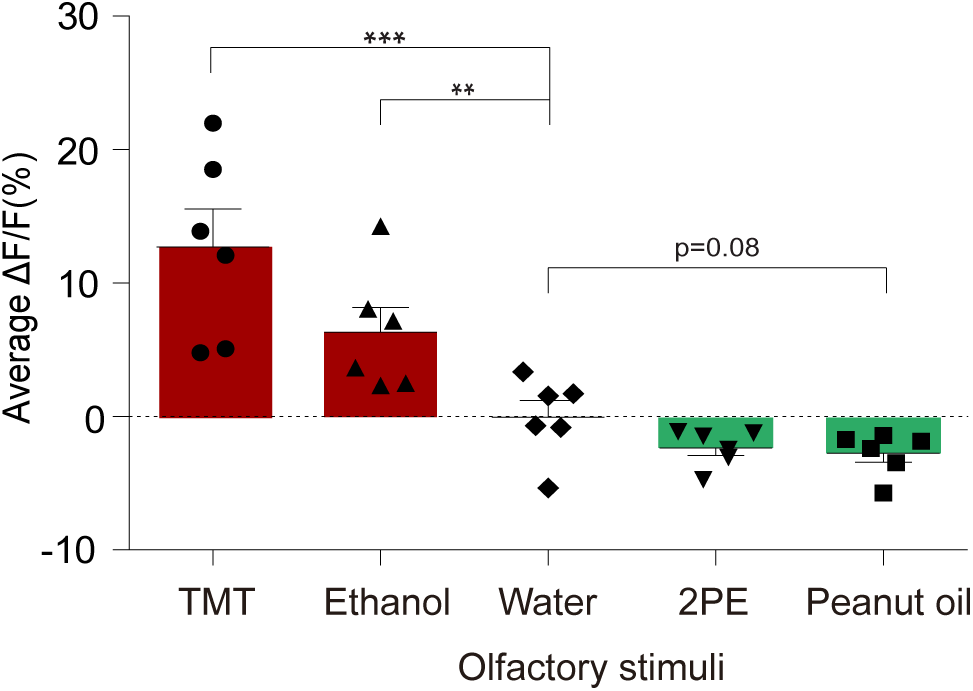
PVN CRF neurons show opposing responses to aversive and attractive odorants. Bar graph summarizing average PVN-CRF^GCaMP^ signal during exposure to each odorant; TMT, Ethanol, water, 2PE (2-phenylethanol), and peanut oil. **p<0.01, ***p<0.001, Data are presented as mean ± s.e.m. Related to Figure 2.

**Supplementary Figure 6.**
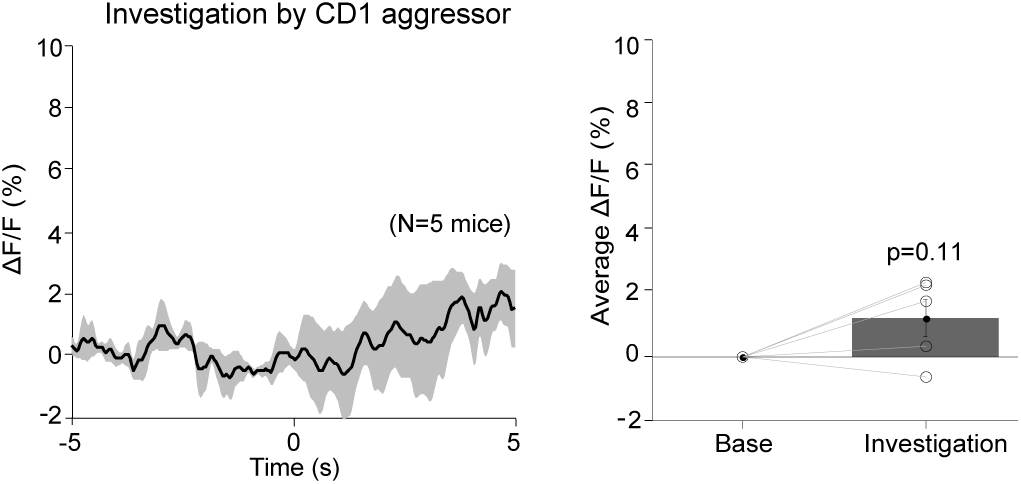
PVN-CRF^GCaMP6^ signal of recorded mice while interacting with CD1 aggressor. A population PETH plot from recorded mice during investigation with aggressive intruders that does not involve in attack (left) and bar graph that measures the changes in ∆F/F of PVN-CRF^GFP^ signal (right) (N=5, p=0.11). Related to Figure 3.

